# Highly diverged novel subunit composition of apicomplexan F-type ATP synthase identified from *Toxoplasma gondii*

**DOI:** 10.1101/321620

**Authors:** Rahul Salunke, Tobias Mourier, Manidipa Banerjee, Arnab Pain, Dhanasekaran Shanmugam

## Abstract

The mitochondrial F-type ATP synthase, a multi-subunit nanomotor, is critical for maintaining cellular ATP levels. In *Toxoplasma gondii* and other apicomplexan parasites, many subunit components, necessary for proper assembly and functioning of this enzyme, appear to be missing. Here, we report the identification of 20 novel subunits of *T. gondii* F-type ATP synthase from mass spectrometry analysis of partially purified monomeric (~600 kDa) and dimeric (>1 MDa) forms of the enzyme. Despite extreme sequence diversification, key F_O_ subunits, a, b and d, can be identified from conserved structural features. Orthologs for these proteins are restricted to apicomplexan, chromerid and dinoflagellate species. Interestingly, their absence in ciliates indicates a major diversion, with respect to subunit composition of this enzyme, within the alveolate clade. Discovery of these highly diversified novel components of the apicomplexan F-type ATP synthase complex could facilitate the development of novel anti-parasitic agents. Structural and functional characterization of this unusual enzyme complex will advance our fundamental understanding of energy metabolism in apicomplexan species.

## Introduction

The F-type ATP synthase is a ubiquitous nanomotor present on the inner membrane of mitochondria, chloroplast and bacterial plasma membrane that catalyzes ATP formation from ADP and Pi [1,2]. In mitochondria, the energy required to drive the nanomotor is harnessed from the proton motive force and associated membrane potential, which are generated by the mitochondrial respiratory complexes [3,4]. The mitochondrial F-type ATP synthase complex consists of two sub-complexes called the F_1_ and F_O_ sectors. The hydrophilic globular F_1_ sector includes three each of α and β subunits, and a central stalk containing one each of γ, δ and ε subunits. The site of ATP formation resides in the catalytic center located in the β subunit [1,2,4,5]. Unlike the F_1_ sector subunit components, which are well conserved across various eukaryotic phyla, many F_O_ sector subunit components are highly diverse [6–9] and are not readily identified based on sequence similarity. The complete set of F_O_ sector subunits have been mapped only in a few species [10,11]. For example, in the yeast *Saccharomyces cerevisiae*, the F_O_ sector is composed of the hydrophobic membrane spanning oligomeric subunit c (10-12 monomers), along with subunits a, b, d, f, 8, h and OSCP (Oligomycin sensitivity confirming protein) [10]. In addition, other accessory subunits such as e, g, i, j and k are also associated with the enzyme. Orthologs of all yeast F_O_ sector subunits, except subunits j and k, are present in the mammalian counterpart (bovine enzyme), which additionally has subunits AGP and MLQ [11].

Numerous biochemical and structural studies have addressed the assembly and interaction between the various subunit components of F-type ATP synthases [1,12]. These studies have provided in-depth insights on the functional coupling of the proton motive force to ATP synthesis. The F_O_ sector subunits c and a interact to form the proton channel, and proton translocation, which occurs along the interface of these two subunits, is coupled to the rotary motion of the c-ring and central stalk of the enzyme [13–16]. The peripheral stalk structure of the enzyme, comprised of F_O_ sector subunits b, d, h (f6 in bovine) and OSCP, has the critical role of holding the α_3_β_3_ catalytic domain in place during the clockwise rotation of the central stalk. These events elicit conformational changes in the α_3_β_3_ catalytic domain, which then results in ATP synthesis [2,4,17,18]. F_O_ sector subunits e and g are known to be involved in dimer assembly of the enzyme complex in yeast [19,20], which appears to facilitate cristae formation by the mitochondrial inner membrane [21–24].

However, despite the importance of the various F_O_ sector subunits, only subunit c and OSCP can be detected from sequence conservation in the vast majority of unicellular eukaryotes, including in the alveolate infrakingdom, which includes the phylum *Apicomplexa* [25–27]. It is unlikely that the missing subunits are totally absent in these species. Plausible scenarios are that they have either diverged in sequence beyond recognition or that a completely new set of proteins have substituted for the missing subunits in these species. This is indeed the case in organisms such as *Euglena gracilis* [9], *Trypanosoma brucei* [7], *Tetrahymena thermophile* [8] and *Chlorophyceae* algae [6]. Hence, our interest was to characterize the complete subunit composition of the F-type ATP synthase from the apicomplexan parasite *Toxoplasma gondii*.

*T. gondii* is an obligate intracellular protozoan parasite, responsible for toxoplasmosis in humans and animals [28]. The parasite completes its life cycle in two host species; the definitive hosts are naïve feline species which support the sexual development of the parasite, and all warm blooded animals are intermediate hosts and support its asexual development [29]. During asexual development, the parasites can reversibly differentiate between fast-growing virulent tachyzoites and slow growing latent bradyzoites [30]. By virtue of their fast-growing nature, tachyzoites are metabolically more active and glycolysis is the primary source for carbon and energy for optimal growth of this parasite [31–33]. However, recent findings have revealed that glucose is not an essential nutrient [34], and parasites survive in the presence of glutamine as an alternative nutrient source [35,36]. In the absence of glucose oxidation *via* glycolysis, oxidative phosphorylation is the only source for bulk cellular ATP. Evidence for an active oxidative phosphorylation in *T. gondii* exists [37–39], and treatment with atovaquone, a potent inhibitor of mitochondrial electron transport chain (mtETC), results in inhibition of ATP synthesis and parasite growth (*EC*_50_ < 10 nM) [38].

Very little is known about the structure and function of the F-type ATP synthase from *T. gondii* and other apicomplexan parasites. Comparative genomics has revealed that, while the five canonical F_1_ sector subunits -α, β, γ, δ, and ε - are readily identified in all apicomplexan parasites based on amino acid sequence conservation, only two F_O_ sector subunits - c and OSCP - can be identified by sequence [25–27]. Characterization of F-type ATP synthase from solubilized mitochondria lysates of the human malaria parasite *Plasmodium falciparum* revealed that the enzyme assembles into monomer and dimer forms [26]. This suggests that a full complement of subunits, typical for the eukaryotic enzyme, is present in *P. falciparum* and likely in other apicomplexan parasites as well.

Here, we have investigated the subunit composition of the enzyme from *T. gondii.* We were able to identify and partially purify the F-type ATP synthase enzyme complex from solubilized mitochondrial preparation of *T. gondii* using blue native PAGE separation, by immuno-precipitation and by chromatographic enrichment. LC-MS analysis of the enzyme preparations revealed the identity of the proteins associated with the *T. gondii* F-type ATP synthase. We have identified 20 novel proteins (of unknown function) as being *bona fide* subunit constituents of *T. gondii* F-type ATP synthase based on consensus from multiple and independent experiments. Phyletic profiling revealed that orthologs for many of these proteins are present in most apicomplexan species, except in those *Cryptosporidium sp* which possess a degenerate mitochondrion devoid of all proteins involved in oxidative phosphorylation [40]. Further, the phyletic profiles of these novel proteins reveal their origin to be ancient, probably even ancestral to alveolate radiation. The identification of these novel protein components of the apicomplexan F-type ATP synthase will facilitate further structural, functional and inhibitor discovery studies on this important enzyme complex.

## Results

### Identification of F-type ATP synthase subunits from T. gondii proteome

Comprehensive *in silico* analysis on the presence and absence of key enzymes involved in mitochondrial metabolism in apicomplexan species has been previously carried out [25,27,41]. These studies have highlighted the missing subunits of the F-type ATP synthase enzyme from these organisms. We have further confirmed these findings from our efforts to identify *T. gondii* orthologs for yeast and bovine F-type ATP synthase subunits. Orthologs for F_1_ sector subunits α (TGME49_204400), β (TGME49_261950), δ (TGME49_226000), ε (TGME49_314820), and γ (TGME49_231910) and F_O_ sector subunits c (TGME49_249720) and OSCP (TGME49_284540) were readily identified from *T. gondii*. This suggested that all the subunits necessary for assembling the catalytic core (α3β3), central stalk (δ, ε and γ) and the subunit c oligomer portions of the F-type ATP synthase (Fig 1A) are encoded by *T. gondii*. However, as reported previously [25,27], we were unable to identify the orthologs for critical F_O_ sector subunits involved in proton translocation and formation of the peripheral stalk structure of the enzyme in *T. gondii* through routine bioinformatic analysis. This appears to be the case in all apicomplexan species and in other taxa within the alveolate infrakingdom, such as the dinoflagellates, ciliates and chromerids (Fig 1B). It is likely that the missing F_O_ subunits are either highly divergent or completely novel. Interestingly, this situation is not exclusive to alveolates, since many other unicellular eukaryotes also appear not to possess the corresponding orthologs for many of the yeast or bovine F_O_ subunits. In fact, novel F-type ATP synthase associated proteins have been previously identified from *Tetrahymena* [8], *Chlamydomonas* [6] and *Trypanosoma* [7]. Therefore, we attempted to purify the F-type ATP synthase enzyme from isolated *T. gondii* mitochondria in native form and identify its protein composition by mass spectrometry.

**Fig 1.**
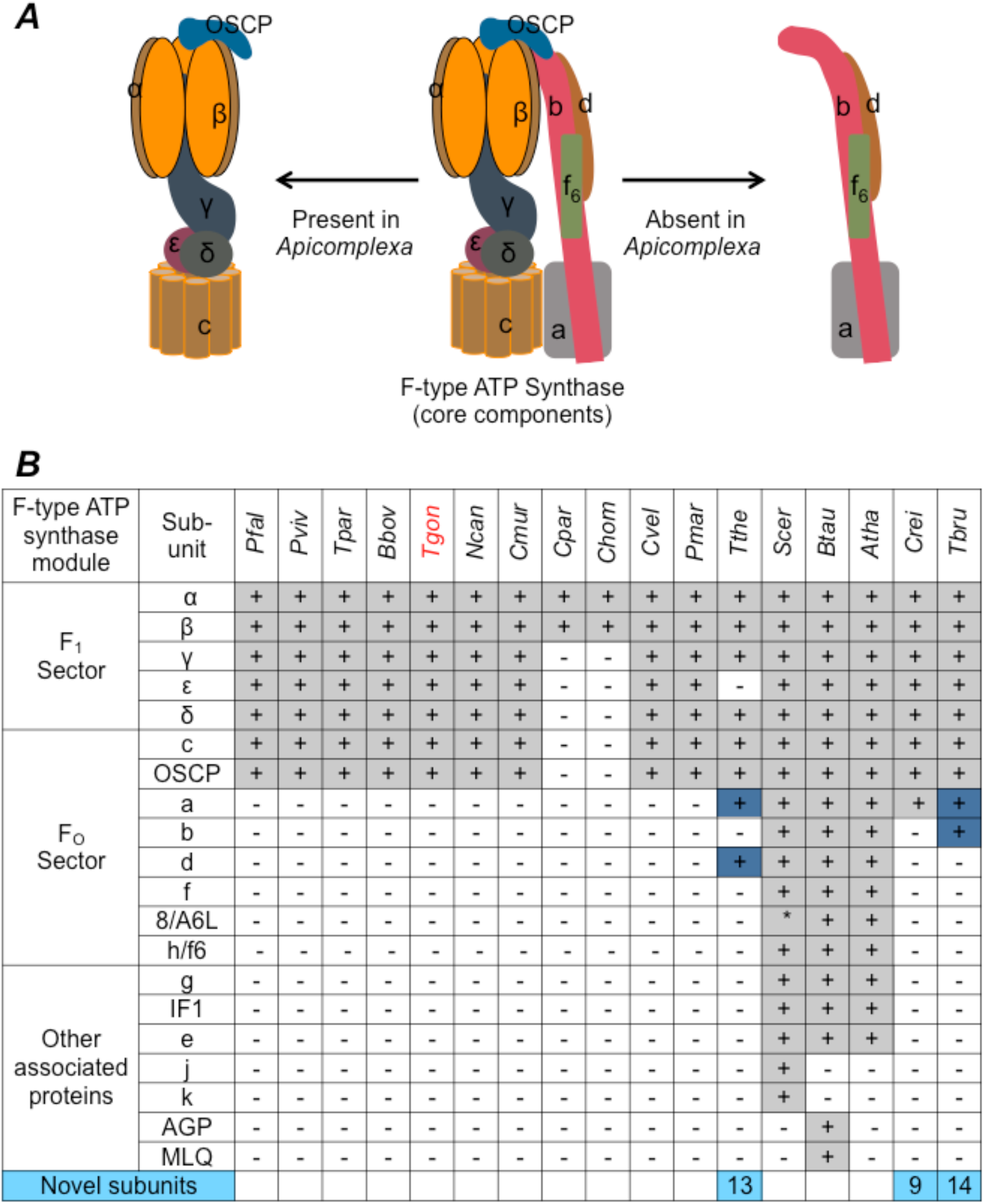
Detail of F-type ATP synthase subunits missing in *T. gondii* and other apicomplexan parasites. (A) Schematic representation of core subunit composition of F-type ATP synthase. The different subunits are color coded and annotated. Middle, the complete set of subunits of the core enzyme; left and right, subunits identified and not identified by *in silico* methods in apicomplexan genome. (B) Table showing the presence/absence of orthologs of yeast and bovine F-type ATP synthase in *Apicomplexa* and other alveolate species. +/-, presence/absence of ortholog; blue, diverged (no clear orthologs) functional equivalent; light blue, experimentally identified species-specific novel subunits (numeric values indicate the total number of such novel subunits). Species names: *Pfal*, *P. falciparum*; *Pviv*, *P. vivax*; *Tpar*, *T. parva*; *Bbov*, *B. bovis*; *Tgon*, *T. gondii*; *Ncan*, *N. caninum*; *Cmur*, *C. muris*; *Cpar*, *C. parvum*; *Chom*, *C. hominis*; *Cvel*, *C. velia*; *Pmar*, *P. marinus*; *Tthe*, *T. thermophila*; *Scer*, *S. cerevisiae*; *Btau*, *B. taurus*; *Atha*, *A. thaliana*; *Crei*, *C. reinhardtii*; *Tbru*, *T. brucei*.

### Generating transgenic parasites expressing YFP-HA tagged TgATP –β and TgATP-OSCP proteins

To facilitate the purification of the *T. gondii* F-type ATP synthase enzyme in native form, we engineered in-frame genomic YFP-HA tags in the 3’ end of the respective genes encoding the F_1_ β and F_O_ OSCP subunits (Fig 2A). The modified genes continued to be expressed under the control of their endogenous promoters. The correct insertion of the tags was confirmed by genomic PCRs (S1 Data) from clonal isolates of the respective transgenic parasites. Expression of the desired YFP-HA tagged proteins, and their mitochondrial localization, were confirmed by Western blotting (Fig 2B) and microscopy (Fig 2C), respectively. We then compared the ability of the parental and transgenic strains of the parasites to maintain total cellular ATP levels in the presence/absence of glucose. Although *T. gondii* tachyzoites are known to prefer glucose as the primary nutrient source, in the absence of glucose they are known to switch to glutaminolysis for carbon and energy supply. In the later case, ATP is obtained by the parasites *via* oxidative phosphorylation, and this can be inhibited using atovaquone [25,42,43]. Like the parental parasites, transgenic parasites expressing YFP-HA tagged F_1_ β and F_O_ OSCP subunits were capable of maintaining cellular ATP levels *via* oxidative phosphorylation in the absence of glucose, which was inhibited by atovaquone (Fig 2D). These results confirm that the modification of either F_1_ β or F_O_ OSCP proteins with YFP-HA tag had no detrimental effect on the function of the enzyme, implicating that the structure of the tagged enzyme remained intact.

**Fig 2.**
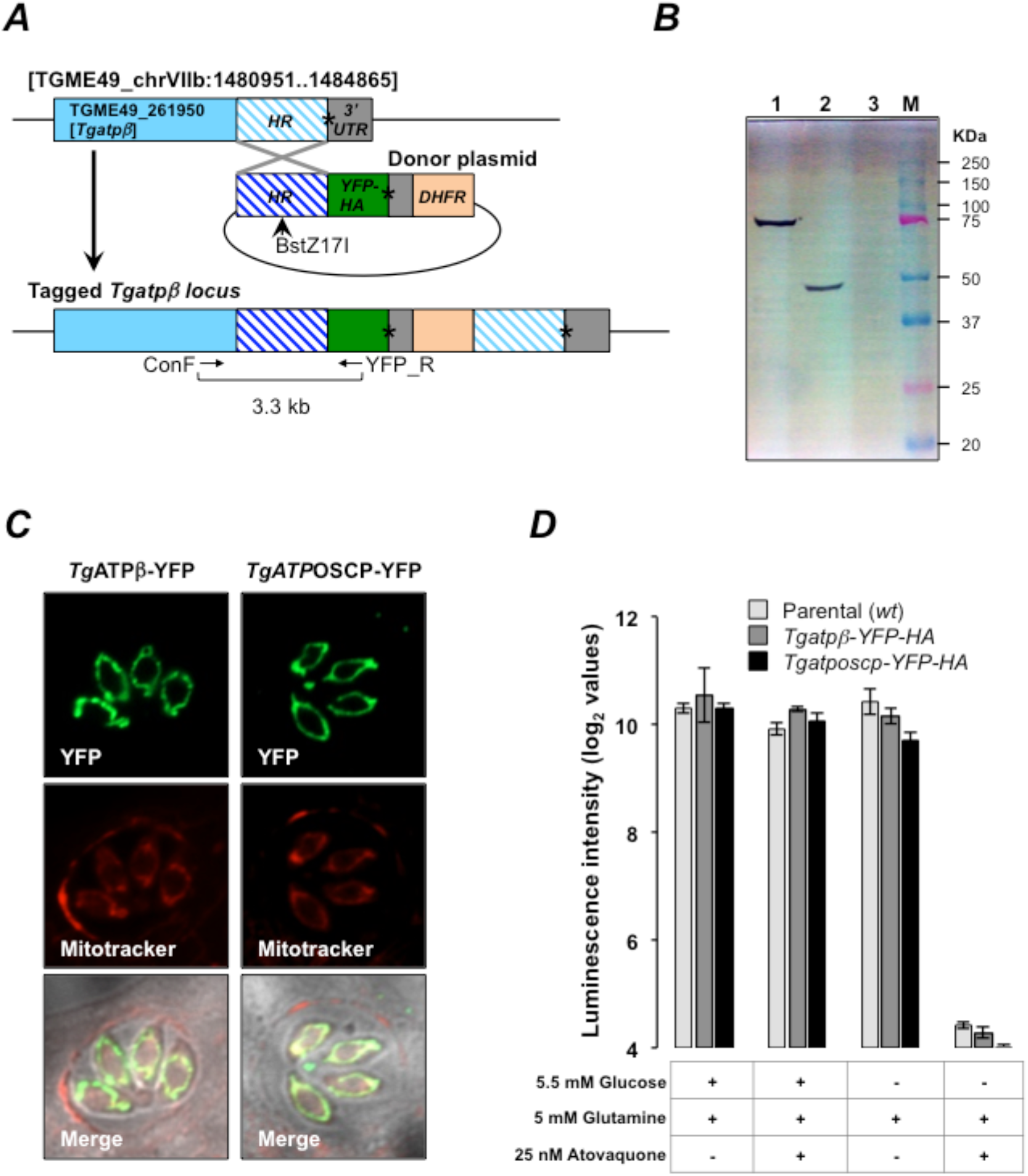
Endogenous tagging of *Tgatpβ* and *Tgatposcp* genes with YFP-HA tags. (A) Schematic representation of the *Tgatpβ* (TGME49_261950) gene locus wing coding region in light blue and 3’ UTR in grey. The hatched region shows the homology region (HR), which was cloned into a donor plasmid (dark blue) upstream of an in-frame coding region for YFP-HA tag. The donor plasmid also includes the DHFR cassette, which confers pyrimethamine resistance. Asterisk denotes the position of stop codon. The donor plasmid was linearized with the restriction enzyme BstZ17I to facilitate single crossover homologous recombination and the resulting modification of the gene locus was confirmed by PCR using the primers ConF and YFP_R. A similar strategy was used to tag the *Tgatposcp* gene locus. (B) and (C) Confirming the expression of the *Tg*ATPβ−YFP-HA and *Tg*ATPOSCP-YFP-HA by SDS-PAGE western blotting (B) and microscopy (C) on respective clonal transgenic lines. Gel lanes in (B): 1, Cell lysate from *Tg*ATPb-YFP-HA transgenic parasites; 2, Cell lysate from *Tg*ATPOSCP-YFP-HA transgenic parasites; 3, Cell lysate from parental parasites; M, Molecular weight size markers. Mitotracker was used to visualize the mitochondrion in (C). (D) Functional validation of mitochondrial ATP synthesis in *wt* parental strain, and *Tgatpβ-YFP-HA* and *Tgatposcp-YFP-HA* expressing transgenic parasites. All three strains exhibited similar response to atovaquone treatment in the presence and absence of glucose, thereby confirming that the YFP-HA tag had no effect on mitochondrial ATP synthesis.

### Identification of monomer and dimer forms of T. gondii F-type ATP synthase by Blue Native Page (BNP) analysis

The fully assembled F-type ATP synthase is known to exist as dimers on the inner mitochondrial membrane and this dimerization is known to influence the characteristic cristae formation by the membrane [21,24]. In case of the human malaria parasite *P. falciparum* F-type ATP synthase, dimer formation has been observed previously [26]. To find out whether the *T. gondii* enzyme can assemble into dimers, we carried out BNP analysis of detergent solubilized mitochondrial preparations. After resolving the samples on a 3% - 12% gradient BNP gel capable of resolving a molecular weight range between 20 kDa to ~1200 kDa (Fig 3A lane M), Coomassie staining revealed prominent bands at the sizes corresponding to dimeric and monomeric forms of the enzyme (Fig 3A lane A). This was further confirmed by Western blotting after BNP separation using α-HA antibodies (Fig 3A lane B), which indicated that the dimer form of the enzyme is more abundant than the monomer form.

**Fig 3.**
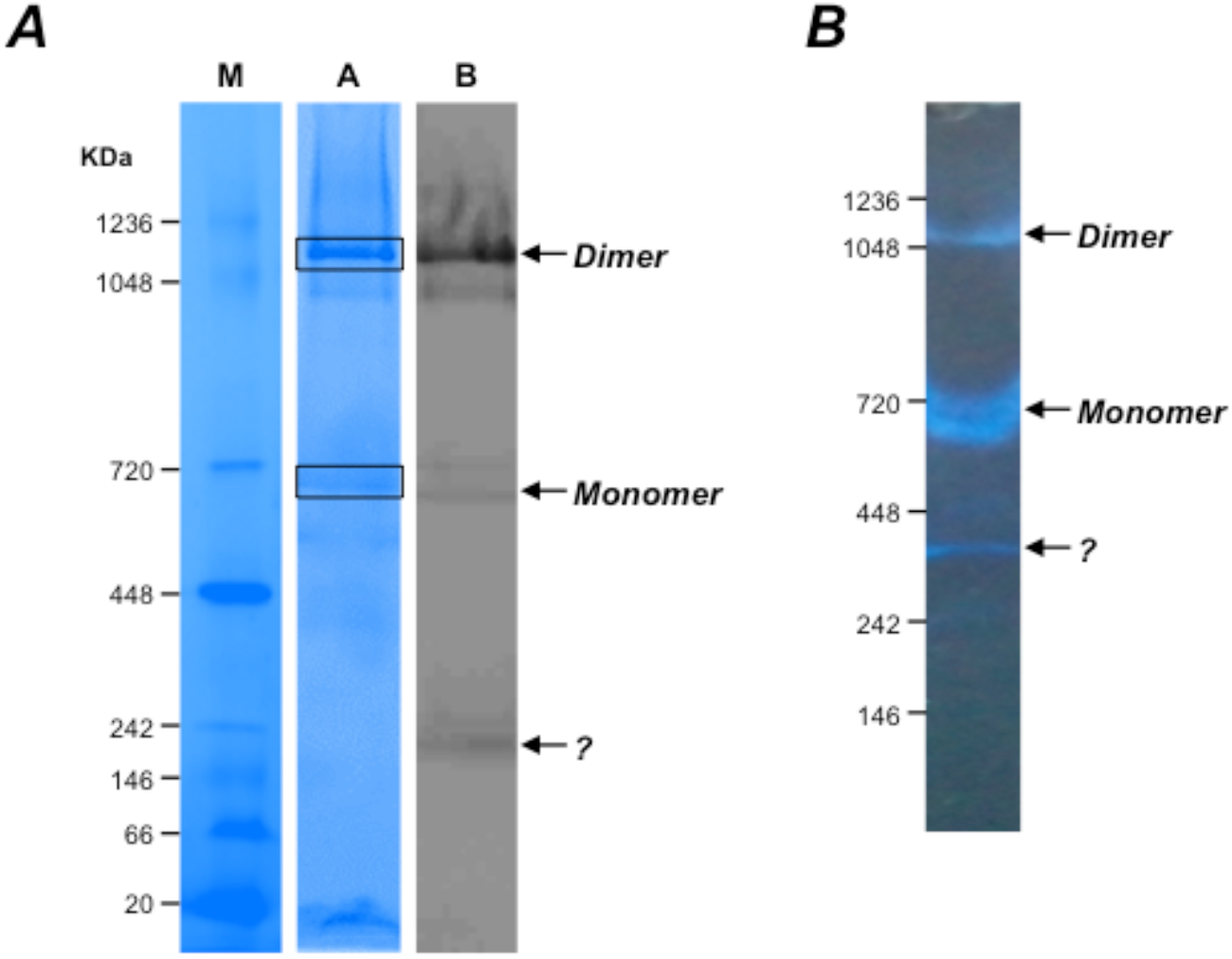
Identifying the dimer and monomer forms of *T. gondii* F-type ATP synthase by BNP analysis. (A) Mitochondria lysates were prepared from tachyzoites stage *Tgatpβ-YFP-HA* transgenic parasites were separated by BNP. Lane M, native molecular weight markers; lane A, Coomassie blue staining of BNP gel; lane B, western blotting of BNP gel using α-HA antibodies. The dimer and monomer forms are indicated by arrows. The boxed region in lane A corresponds to the excised gel piece, which was processed for LC-MS/MS analysis. (B) In-gel ATPase activity assays following BNP separation confirms that the dimer and monomer forms of F-type ATP synthase are functionally intact.

Next, we carried out in-gel activity assays, to detect the ATPase activity associated with the isolated enzyme. Results from these assays confirmed that the ATPase activity was intact, for both dimer and monomer forms of the enzyme, after BNP separation (Fig 3B). We found that, although the dimer form of the enzyme was more abundant, its ATPase activity was less than that of the monomer form. This is supported by the fact that F_1_ ATPase activity is inhibited *in vivo* to minimize the risk of ATP hydrolysis and favor ATP synthesis [44]. Since the in-gel activity assay is an ATP hydrolysis assay, the dimer has less of this activity. Based on BNP mobility, we deduce the size of the dimer form of the enzyme to be between 1 – 1.2 MDa and that of the monomer form as ~600 kDa. This is in agreement with what has been reported previously for *P. falciparum* [26], *S. cerevisiae* [19], and bovine [45] enzymes, and suggests that a full complement of F_O_ subunits is present in *T. gondii* F-type ATP synthase. Therefore, we excised the regions in the BNP corresponding to the dimer and monomer forms of the enzyme and proceeded to identify the proteins in the gel band by LC-MS analysis (Fig 3A).

### Identification of T. gondii F-type ATP synthase associated proteins by LC-MS/MS proteomics

A standard protocol (see methods section) was followed for in-gel LC MS/MS analysis of the gel bands corresponding to the dimer and monomer forms of the enzyme identified by BNP. This was done on samples prepared from both *Tgatp−β−YFP-HA* and *Tgatp-OSCP*−*YFP-HA* transgenic parasites. Since the dimer form of the enzyme was more abundant than the monomer form after BNP separation, we had better success with LC-MS/MS analysis of the gel band corresponding to the dimer form of the enzyme. A total of 96 proteins were identified with high confidence from consensus data obtained from multiple in-gel LC MS/MS analysis (Fig 5). Proteins corresponding to all F_1_ subunits and F_O_ OSCP were detected, while FO subunit c was not detected in any of the experiments. In addition, many proteins of unknown function were also detected. However, due to co-migration of several non-specific proteins, such as myosin, in the area corresponding to the gel band processed for LC MS/MS analysis, it is likely that some of these proteins of unknown function are not *bona*-*fide* subunits of the F-type ATP synthase

In order to specifically identify the protein subunits of F-type ATP synthase, we resorted to partially purify the enzyme before LC MS/MS analysis by chromatographic separation (Fig 4A, B) and immunoprecipitation (IP; using the α-HA antibody). In contrast to BNP analysis, after size exclusion chromatography we observed that the dimer form of the enzyme was less abundant than the monomer form (Fig 4B). This is likely due to stability issues with the dimer, which might progressively fall apart as monomers during the process of chromatographic separation. The fractions corresponding to the dimer and monomer forms of the enzyme were pooled and concentrated separately before processing for LC-MS/MS analysis. A total of 64 proteins were detected (Fig 5A) from the fractions corresponding to monomer form of the enzyme and we did not get reliable data from the dimer fractions owing to very low protein concentration.

**Fig 4.**
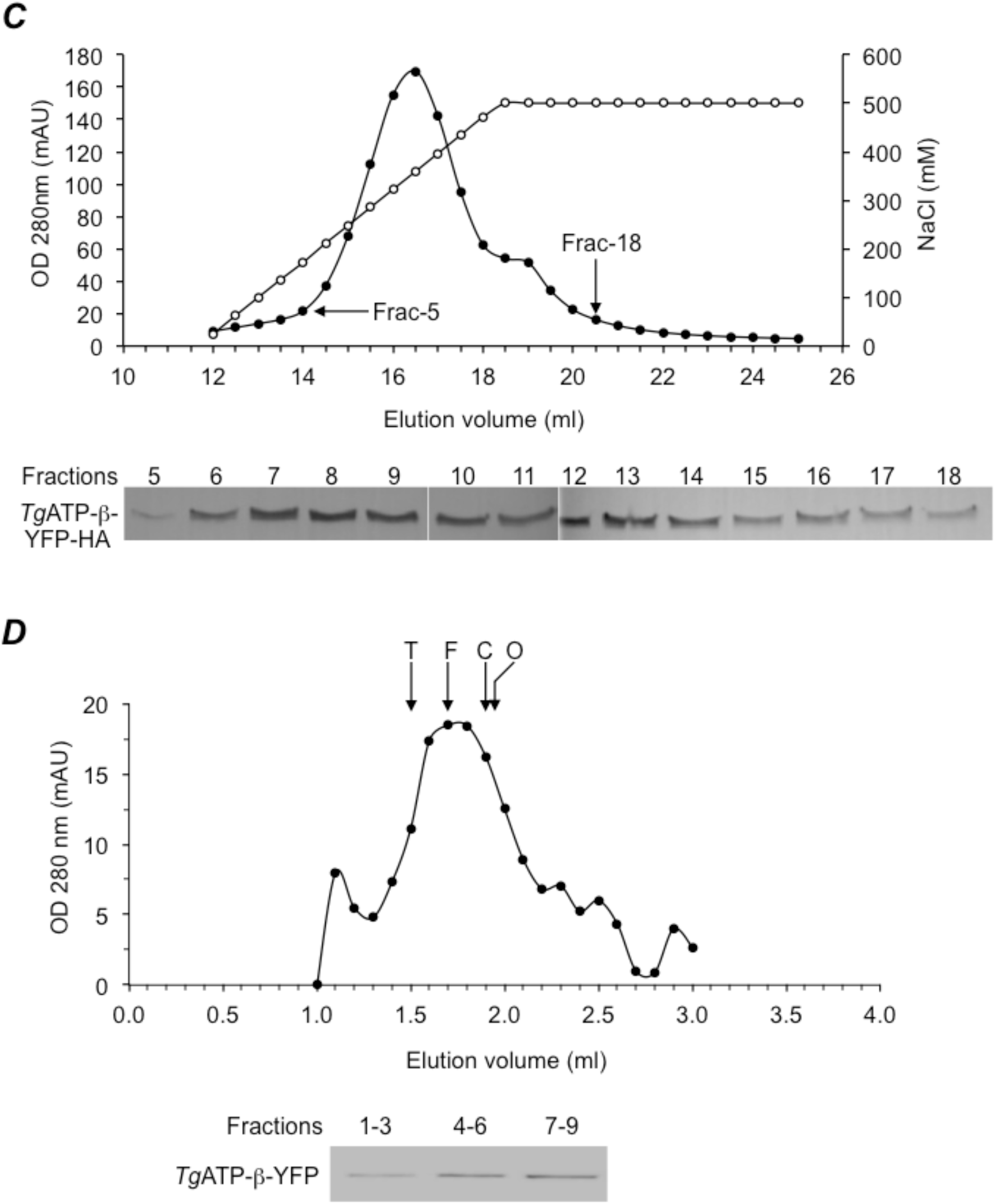
Partial purification of dimer and monomer forms of *T. gondii* F-type ATP synthase by chromatography. (A) Ion exchange (DEAE sepharose) separation of mitochondrial lysates prepared from *Tgatpβ-YFP-HA* transgenic parasites. Absorbance at 280 nm (filled circles) and NaCl concentration (open circles) are plotted for each fraction. Fractions 5 and 18 are marked with arrows. Bottom panel shows SDS-PAGE western blotting for fractions 5-18 to find out the elution profile of *Tg*ATPβ-YFP-HA. (B) Size exclusion profile of the pooled fractions from ion exchange chromatography. The Absorbance at 280 nm is plotted for each fraction. The size exclusion column was calibrated with the following native markers; thyroglobulin (T, 660 kDa), ferritin (F, 440 kDa), conalbumin (C, 75 kDa), Ovalbumin (O, 45 kDa). Peakelution volume for each marker is indicated by arrow. Fractions 1-3, 4-6 and 7-9 were pooled, concentrated and subject to SDS-PAGE western blotting to detect *Tg*ATP-β-YFP-HA as shown in bottom panel.

**Fig 5.**
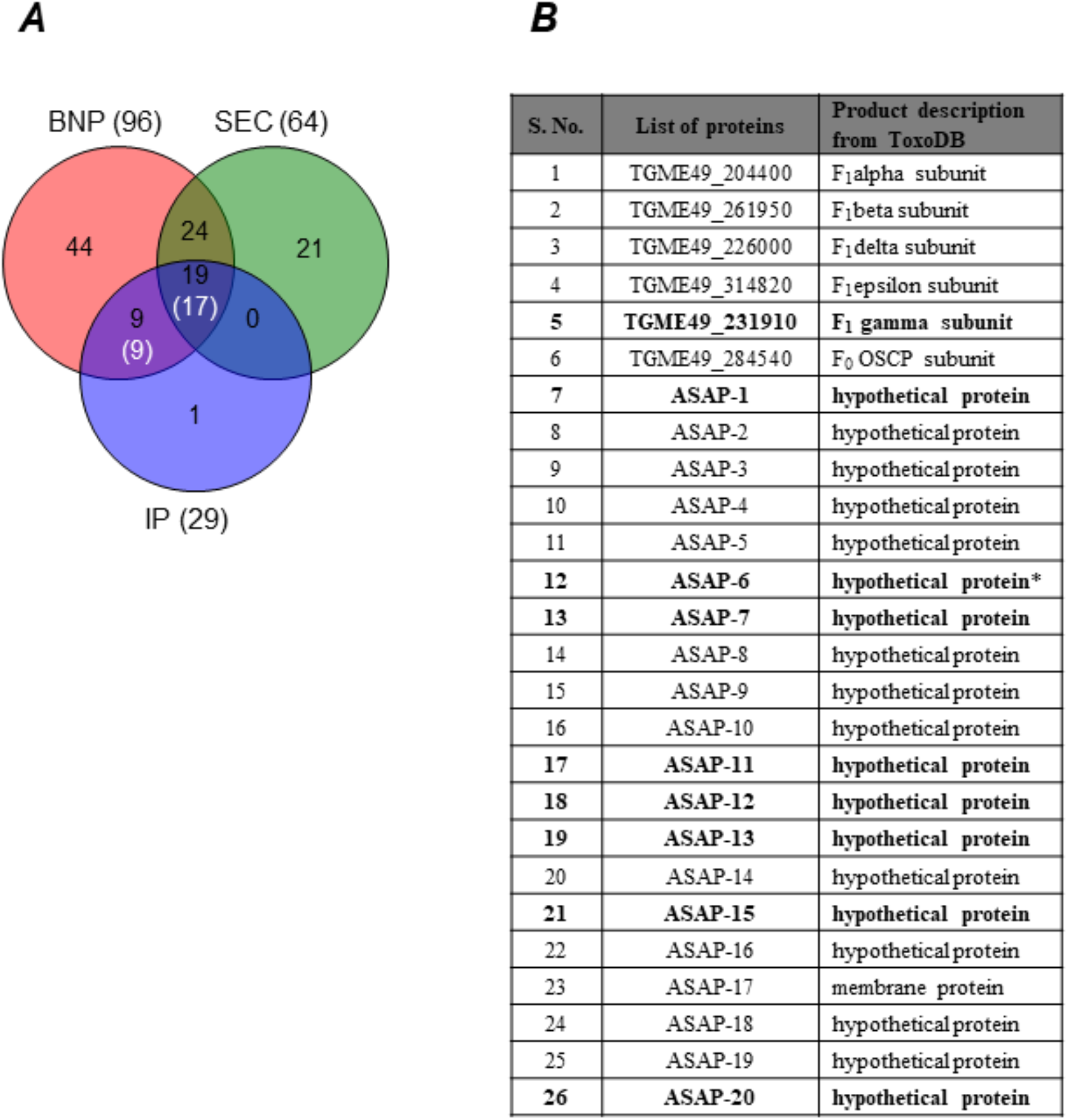
Identification of novel *T. gondii* F-type ATP synthase subunits from LC-MS/MS analysis. (A) The Venn diagram shows shared identification of proteins following blue native page (BNP), size exclusion chromatography (SEC) and immunoprecipitation (IP) sample preparation. Total proteins identified in each technique is given within brackets outside the Venn diagram. The numbers shown in white font and within brackets are the final set of proteins assigned as subunit components of *T. gondii* F-type ATP synthase. (B) List of all ASAPs identified in this work by mass spectrometry analysis. The entries shown in bold indicate that the protein was detect with high confidence from only BNP and IP samples. Asterisk indicates that the corresponding peptides were detected in only one of the replicate runs for SEC sample. BNP data is a combination of experiments done with both *Tg*ATPβ-YFP-HA and *Tg*ATPoscp-YFP-HA expressing transgenic parasites. SEC and IP data are from *Tgatpβ-YFP-HA* and *Tgatposcp-YFP-HA* transgenic parasites respectively.

LC-MS/MS analysis on the enzyme enriched by IP resulted in the identification of a total of 29 proteins, of which 28 were also detected from either BNP or chromatography samples (Fig 5A). It is notable that 19 proteins, including most F_1_ subunits and FO OSCP, were detected in samples prepared by all three methods. We detected very few non-specific proteins from the IP samples, i.e., functionally not related to F-type ATP synthase (3 out of 29). Out of the remaining 26 proteins, 5 were known F_1_ subunits (α, β, δ, ε, γ), one was F_O_ OSCP, and the remaining 20 were proteins of unknown function. It should be noted that we were unable to identify F_O_ subunit c in any of our LC-MS/MS analysis, probably owing to its highly hydrophobic nature. Based on the consensus of proteins identified by the three different approaches, the F-type ATP synthase from *T. gondii* appears to be comprised of at least 27 protein subunits. We have coined the termed ‘ATP synthase Associated Proteins’ (ASAP) to refer to the 20 novel subunit components of the enzyme identified in this study. A complete list of all *T. gondii* F-type ATP synthase subunits identified by mass spectrometry, along with the experiments from which they were detected, is given in the table in Fig 5B.

### Identification of F_O_ subunits a, b and d

As shown in Fig 1, in addition to subunits c and OSCP, the F_O_ sector consists of at least 6 other subunits in yeast, mammalian and plant enzymes. Subunit a is essential for proton conductance, which it facilitates along with subunit c [13–15]. Subunit b forms the core of the stator structure, which is essential for holding the catalytic α_3_β_3_ structure in place during the rotary motion of the central stalk [18]. Subunit d is part of the stator structure. We attempted to find out which of the ASAPs correspond to these three subunits in *T. gondii* F-type ATP synthase. Since none of the ASAPs had any sequence identity to the yeast F_O_ a, b and d subunits, we resorted to conserved structure based identification using pairwise comparison of profile hidden Markov models as implemented in HHPred [46,47]. For this analysis we used the corresponding tool available from web based MPI bioinformatics toolkit (*toolkit.tuebingen.mpg.de*). We analyzed each of the ASAPs using this tool and successfully identified hits for F_O_ subunits a, b and d based on previously known structures for these proteins from other species. The ASAP-1 was identified as F_O_ subunits a (Fig 6A), and we were also able to generate sequence alignments for the C-terminal domain of this protein which showed the conservation of key amino acid residues arginine (important for proton translocation) and glutamine (yellow highlight in Fig 6B). Similarly, searches with ASAP-2 and ASAP-3 came up with hits for F_O_ subunits b and d, respectively (S2 Data and S3 Data).

**Fig 6.**
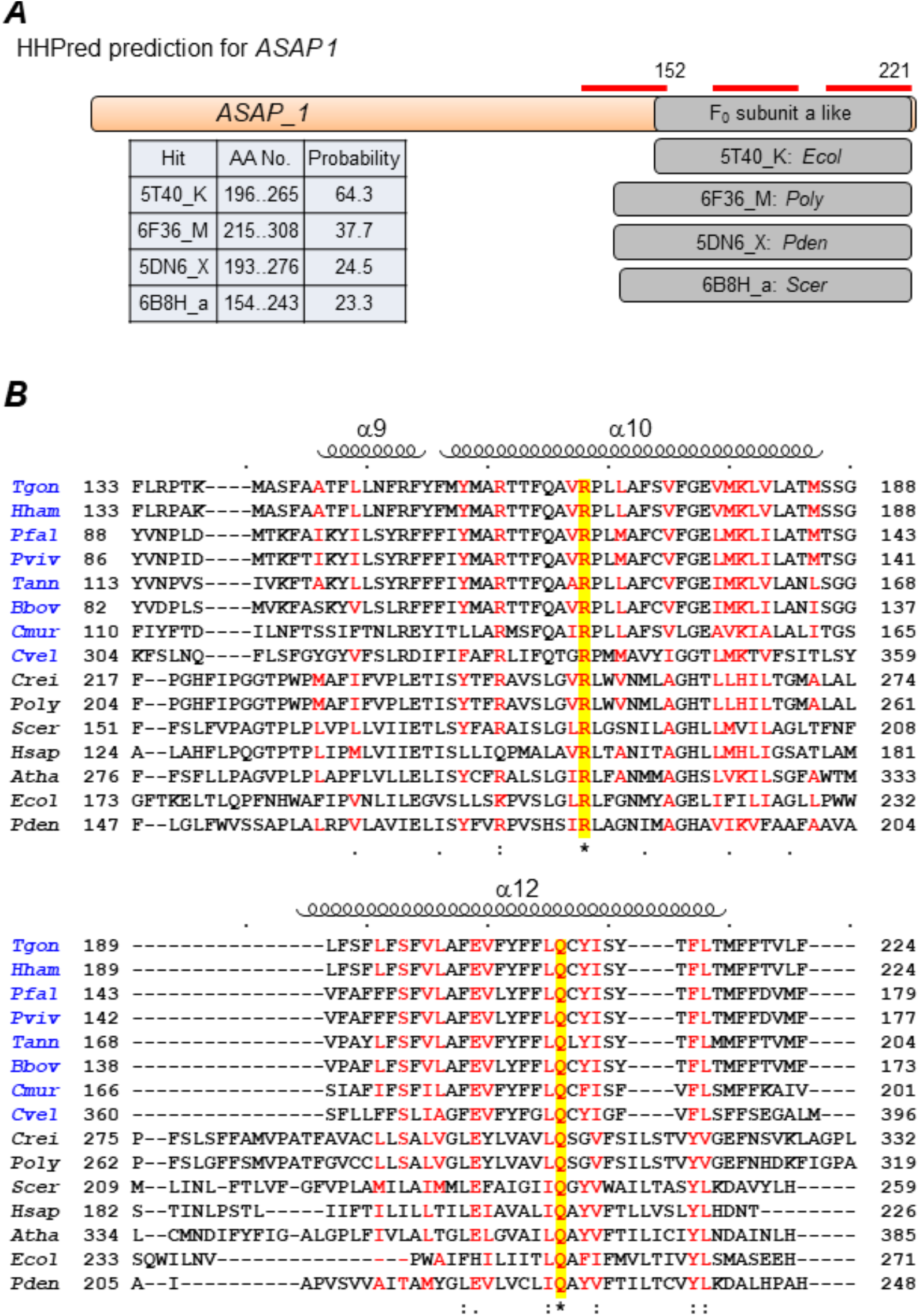
HHPred identification of novel F_O_ subunit a based on conserved structural features. (A) Representation of the pairwise sequence alignments generated by HHPred [46,47] for the putative F_O_ subunit a from *T. gondii* and four other F_O_ subunit a proteins for which the structure is known. The table provides details of the amino acid length and a probability score for the prediction from the hit alignments. The red lines indicated the 3 transmembrane domains present in the *T. gondii* protein. (B) Protein sequence alignments and secondary structure information were made using the Clustal Omega [72] and ESPript [73] softwares, and only the C-terminal portion of respective proteins is shown. The names of alveolate species are highlighted in blue. For the non-alveolate species included in the alignment, the F_O_ subunit a is either readily identified from sequence (*Scer, Hsap, Atha, Ecol* and *Pten*), or has been experimentally determined (*Crei* and *Poly*). Positions with similar amino acids are highlighted in red in the alignment. The arginine and glutamine residues, highlighted in yellow, are conserved in all species and important for function. The helices (α9, α10 and α12) shown are from the structure of *Poly* F-type ATP synthase. Species names: *Tgon*, *T. gondii*; *Hham*, *H. hammondi*; *Pfal*, *P. falciparum*; *Pviv*, *P. vivax*; *Tann*, *T. annulata*; *Bbov*, *B. bovis*; *Cmur*, *C. muris*; *Cvel*, *C. velia*; *Crei*, *C. reinhardtii*; *Poly*, *Polytomella sp.*; *Scer*, *S. cerevisiae*; *Hsap*, *H. sapiens*; *Atha*, *A. thaliana*; *Ecol*, *E. coli*; *Pden*, *P. denitrificans*.

### Mitochondrial localization of ASAPs

Confirming the mitochondrial localization for the ASAPs is essential for validating their subunit membership in *T. gondii* F-type ATP synthase. We first predicted the presence of signal sequence for mitochondrial targeting using the Mitoprot tool [48] in twelve ASAPs (Fig 7). Five ASAPs, including ASAP-2 (F_O_ subunit b like protein), were previously shown to localize in the mitochondrion [49], and we experimentally confirmed the mitochondrial localization for another four (S4 Data). Moreover, unpublished results from Hyper-LOPIT analysis [50] of *T. gondii* tachyzoite proteome in an independent study (personal communication, Dr. Ross Waller, Cambridge University, UK), confirmed that all ASAP proteins (except ASAP-1 and TGME49_249720, for which no data were available) are localized to the mitochondria (Fig 7).

**Fig 7.**
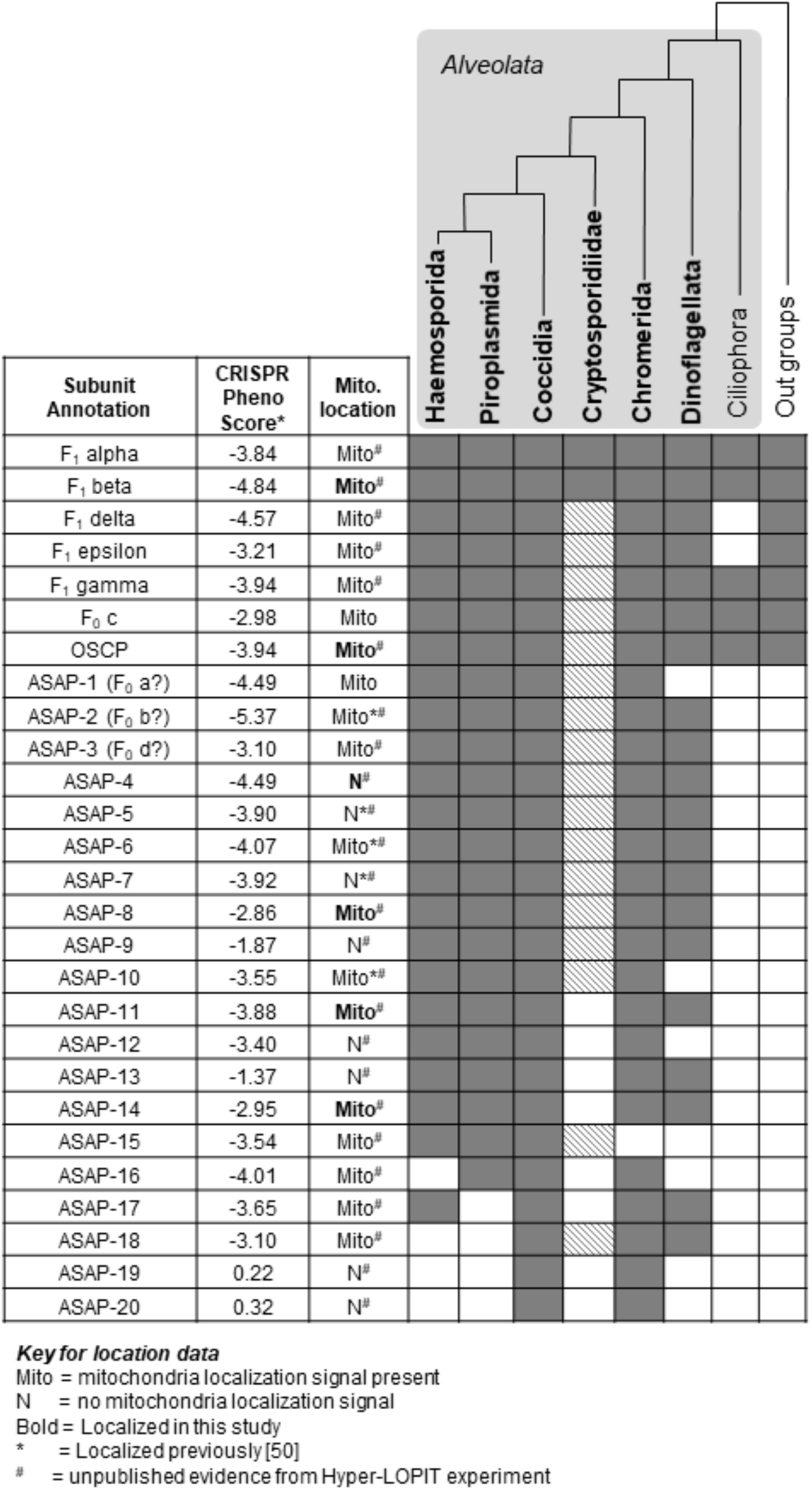
Phylogenetic profile of the alveolate infrakingdom for all subunits mapped to *T. gondii* F-type ATP synthase. Alveolate clades are highlighted with a grey background and their expected phylogenetic relationship is indicated by a tree structure above. Grey and white boxes indicate the presence and absence of the corresponding ortholog, respectively. The hatched boxes represent the presence of the ortholog in *C. muris* only and absence in *C. parvum* and *C. hominis*. The table on the left lists the gene ID for all ASAPs, along with the annotation, essentiality phenotypes (phenotype score from CRISPR/Cas9 knockout [49]), and protein localization experiments from an independent unpublished study (see text in Materials and Methods for details). The key for localization annotation is given below the table.

### Phylogenetic analysis reveals conserved F-type ATP synthase subunit composition in three major alveolate taxons

From our ortholog identification protocol, and from previous studies [27,41], it was apparent that the F-type ATP synthase subunits missing from *T. gondii* were also missing from all other species grouped within the *Alveolata* infrakingdom. In fact, the novel subunit components identified from *Tetrahymena* (*Alveolata*; *Ciliophora*) enzyme were found to be unique to ciliates and not conserved in other alveolate organisms [8]. Therefore, we were interested in finding out whether the novel ASAPs identified in this study are unique to *T. gondii* F-type ATP synthase. Interestingly, we were able to identify orthologs for many of the ASAPs in three major alveolate taxons – *Apicomplexa*, *Chromerida* and *Dinoflagellata* (Fig 7). Out of the 20 ASAPs, 15 were conserved in all apicomplexan clades, except in case of *Cryptosporidium*, where only *C. muris* contained orthologs for 10 of these proteins. A few ASAPs were unique to the *Coccidian* clade. More importantly, all ASAPs, except one, were conserved in *Chromerida*, and at least 9 and 12 ASAPs were also conserved in *Symbiodinium* and *Perkinsus* respectively.

To obtain further insights on the evolutionary origin of these proteins, we generated neighbor-joining phylogenetic trees for ortholog sequences of all F_1_ and F_O_ subunits from representative species of *Haemosporida*, *Piroplasmida*, *Coccidia*, *Cryptosporidiidae*, *Chromerida, Dinoflagellata* and *Ciliophora* (Fig 7 and S5 Data). *Stramenopile* (*Ectocarpus siliculosis*, *Phaeodactylum tricornumtum*, *Pythium ultimum*, *Thalassiosira pseudonana*) and *Plantae* (*Chlamydomonas reinhardtii* and *Arabidopsis thaliana*) species were included as outgroups while constructing the phylogenetic trees for the highly conserved F_1_ subunits. Even though the topology of most trees did not reflect the expected evolutionary relationship between the included species, monophyletic grouping was observed in general at the taxon level, and importantly, this was evident for the conserved F_1_ / F_O_ subunits, as well as the novel ASAPs (S5 Data). Thus, the evolutionary origin of the newly identified highly divergent ASAPs in *Apicomplexa*, *Chromerida* and *Dinoflagellata* clades appears to be ancient.

### Essentiality and gene co-expression analysis of ASAPs from T. gondii and P. falciparum

Due to the functional importance of the F-type ATP synthase enzyme, it is reasonable to expect that the enzyme would be essential in *T. gondii*. This is indeed the case, and all known subunits and ASAPs (except ASAP-19 and 20; Fig 7) of the enzyme were found to be essential in a previous study [49]. In order to obtain further independent evidence for ASAPs as *bona fide* subunits of F-type ATP synthase, we carried out transcriptome co-expression correlation analysis using publicly available gene expression datasets for *T. gondii* and *P. falciparum* [51,52]. We assumed that the ASAPs interacting with one other would show significant pair-wise correlation profiles in their expression levels. Strikingly, we found very high correlation of gene co-expression profiles among the F-type ATP synthase subunits, in comparison to co-expression with other unrelated gene pairs, in both *T. gondii* and *P. falciparum* transcriptome datasets (Fig 8). This finding further supports the fact that the novel ASAPs are indeed co-expressed together and are likely *bona fide* subunits of the unusual F-type ATP synthase enzyme from *T. gondii*.

**Fig 8.**
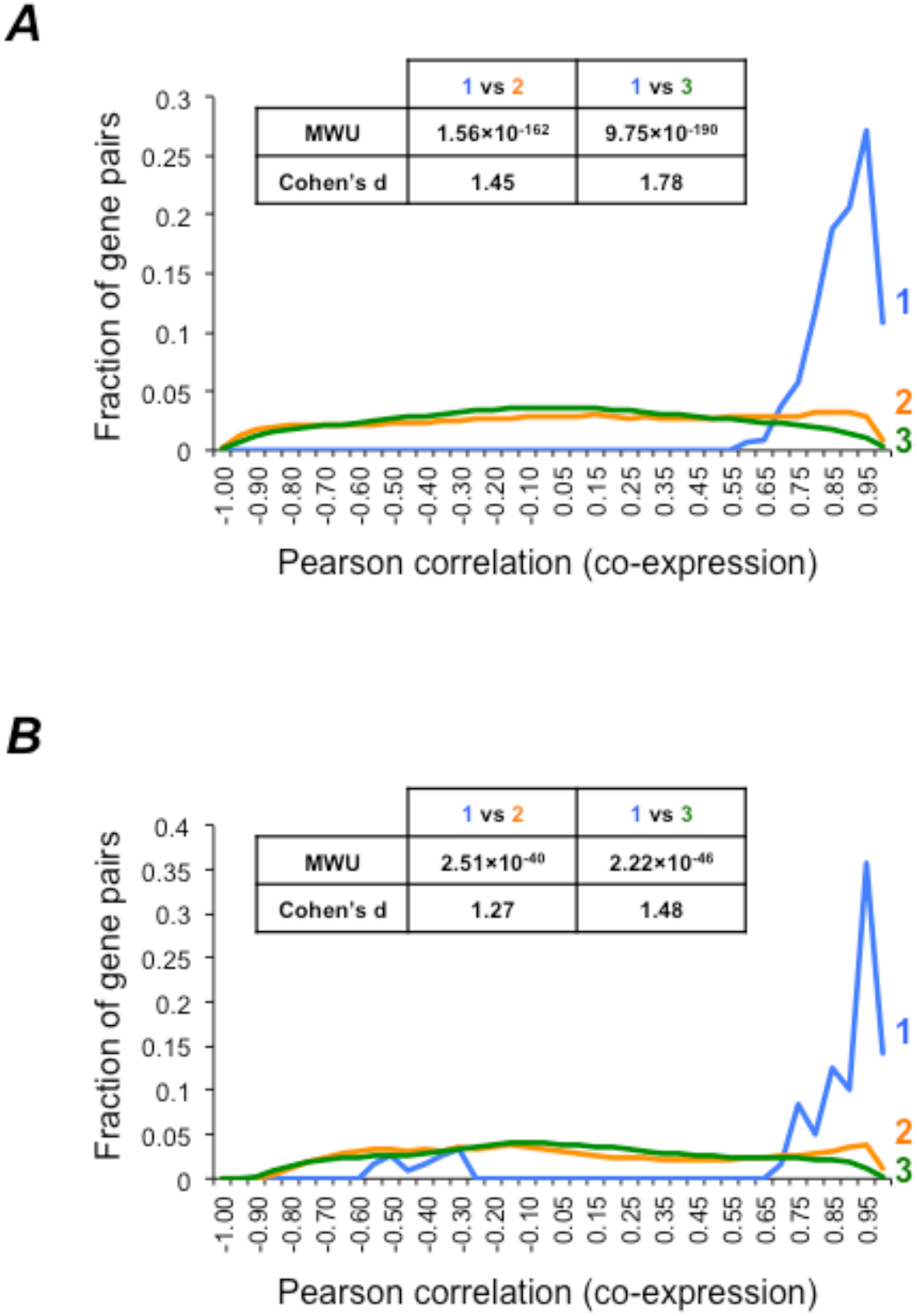
Gene co-expression analysis for *T. gondii* and *P. falciparum* F-type ATP synthase subunits. The distribution of co-expression values - as measured by the Pearson correlation coefficients – are plotted for the three gene pairs categories, as shown for *T. gondii* (A) and *P. falciparum* (B). The statistical support from Mann-whitney U p-values and Cohen’s d values are shown in the table within each plot. Blue (1), co-expression correlation between ATP synthase genes; Orange (2), co-expression correlation between ATP synthase genes and non-ATP synthase genes; Green (3), co-expression correlation between non-ATP synthase genes. MwU, Mann-whitney U p-values; Cohen’s d, Cohen’s d values.

## Discussion

Mitochondrial oxidative phosphorylation is an important source of ATP in most eukaryotic organisms, and is facilitated by the multimeric F-type ATP synthase enzyme complex [1,2,5]. The enzyme consists of two functionally distinct F_1_ and F_O_ sectors, which act in concert to convert the electrochemical energy into mechanical energy to facilitate ATP synthesis. F_O_ sector subunits a and c come together to form the proton translocation channel across the inner mitochondrial membrane. Subunit c, which forms a membrane embedded oligomeric ring, converts the proton gradient energy into mechanical energy *via* subunit rotation. The F_1_ sector α_3_β_3_ catalytic core is connected to the rotary motor formed by F_O_ subunit c ring *via* the rotating central stalk structure comprised of the γ, δ and ε F_1_ subunits [1,2,5]. The stator structure, comprised primarily of the F_O_ subunits b, d, h and OSCP, along with other accessory subunits helps in holding the α_3_β_3_ catalytic core in place while the central stalk rotates [17]. The asymmetric structure of the γ subunit of the rotating central stalk induces conformational changes in the α_3_β_3_ catalytic core, which facilitates ATP synthesis. Thus all subunit components and structural features of this unique enzyme are critical for efficient ATP synthesis.

In addition to the core subunit composition described, other accessory proteins participate in assembling the V shaped dimer form of the enzyme complex [19]. The paired arrangement of the F_1_ complex (spaced by 12 nm) was observed in *Paramecium* inner mitochondrial membrane by freeze-fracture and deep-etching studies [53]. Subsequent studies on yeast, bovine and other species confirmed the dimeric arrangement of the F-type ATP synthase complex by BNP, atomic force microscopy and cryo-electron microscopy [19,54,55]. In fact, the dimer form of the enzyme appears to facilitate cristae formation by the inner mitochondrial membrane [21,22,24] and is implicated in maintenance of the mitochondrial membrane potential [56]. However, the identity of the proteins responsible for dimer formation has not been ascertained in most species. In case of the yeast enzyme, subunits e, g, k were identified as dimer specific components, and studies with genetic mutants revealed that subunits e and g are essential in dimer formation [19,20]. Orthologs for these proteins are present in mammalian species as well.

BNP and size exclusion chromatography studies on purified yeast F-type ATP synthase revealed that the average molecular size of the dimer and monomer form of the enzyme is around 1 MDa and between 500 – 600 kDa, respectively [19]. Given the complex structure and assembly of the enzyme, the expected number of protein subunits in an intact dimer is around twenty, based on yeast and mammalian enzyme compositions [10,11]. Interestingly, while the F_1_ subunits are well conserved across various taxa, the F_O_ subunits, except c and OSCP, appear to be poorly conserved and in many instances are not readily identified from sequence. This is especially true for a variety of unicellular eukaryotes, including many free living and parasitic protists, as evident from KEGG data (*kegg.jp*). Comparative genomics studies revealed that orthologs for yeast and mammalian F_O_ subunits, except c and OSCP, are missing in the entire apicomplexan phylum [25–27]. In fact, along with *Apicomplexa*, this appears to be the case for all other taxa belonging to the alveolata infrakingdom. A study on the subunit composition for F-type ATP synthase from the ciliate *Tetrahymena thermophila*, an alveolate species, resulted in the identification of 13 novel proteins [8]. Surprisingly, these proteins were unique to ciliates, indicating the possible existence of unique set of F_O_ components in other alveolate species as well.

The *P. falciparum* enzyme was identified in dimer form with a molecular size of > 1 MDa, confirming the presence of novel F_O_ subunits [26]. This study also revealed that the enzyme is essential in *P. falciparum*. However, in case of *P. berghei*, F_1_ subunit β was found to be essential for development in mosquito, but not for survival of blood stage parasites in mammalian host [57]. Studies with metabolic mutants in *T. gondii* have revealed that mitochondrial oxidative phosphorylation is an important source of ATP, especially during glucose deprivation and in the absence of a glycolytic flux [34,35]. Mitochondrial ATP synthesis is also likely essential for formation and maintenance of tissue cyst forms of *T. gondii* in infected hosts. Despite the importance of F-type ATP synthase in apicomplexan parasites, not much is known about its subunit composition, structure and function in these parasites.

In this study, we have successfully identified the subunit constituents of F-type ATP synthase from *T. gondii*, a model apicomplexan parasite. Intact monomer and dimer forms of the enzyme can be identified by BNP analysis from detergent solubilized parasite mitochondria preparations. We have identified 20 novel proteins (ASAPs) as subunit constituents of the *T. gondii* F-type ATP synthase by LC-MS/MS analysis of partially purified enzyme using BNP, IP and chromatographic techniques. While some of these proteins could be counterparts of the missing F_O_ subunits, others are most likely accessory proteins which may be involved in dimer formation. Importantly, we were able to identify putative F_O_ subunits a (ASAP-1), b (ASAP-2) and d (ASAP-3), based on conserved structural features. Although the F_O_ subunit a from yeast is known to have six transmembrane domains, the *T. gondii* protein has only three. Nevertheless, the critical arginine residue, which is important for proton translocation [14], appears to be conserved (Fig 6). The hallmark feature of F_O_ subunit b from yeast and other species is the presence of an extended helix that spans the distance between the membrane and the α_3_β_3_ catalytic core. Although there was very poor sequence conservation, very high structural similarity (>97% probability in HHPred assignment) can be found between yeast and *T. gondii* F_O_ subunit b proteins. F_O_ subunit d has no transmembrane domain and is known to interact with F_O_ subunit b *via* parallel/anti-parallel coiled coil helical domains [2]. The putative *T. gondii* F_O_ subunit d was highly similar in secondary structure features over the conserved regions with the bovine enzyme. Further follow up work on the 3D structure of the enzyme is needed to verify these findings.

Interestingly, all F_1_ subunits, and F_O_ c and OSCP subunits, along with 18 ASAPs, were found to be essential in *T. gondii*, in a genome wide CRISPR/Cas9 mediated gene knockout screen [49] (Fig 7). In the same study, some ASAPs were shown to be localized in the mitochondrion. We interrogated the mitochondrial localization for all subunits of *T. gondii* F-type ATP synthase by first considering evidence from *in silico* prediction of mitochondrial localization signals, followed by experimental localization of selected ASAPs (Fig 7). We also obtained confirmation for mitochondrial localization of all but two ASAPs from Hyper-LOPIT analysis of tachyzoite stage *T. gondii* (unpublished data from an independent study, Dr. Ross Waller, University of Cambridge, UK; personal communication). Further, we found that the transcript co-expression for the subunits of F-type ATP synthase was highly correlated, independently in both *T. gondii* and *P. falciparum* transcriptome datasets (Fig 8), thus providing additional evidence for functional interaction between these proteins across apicomplexan parasite taxa. In summary, evidence from gene essentiality, subcellular localization and transcript co-expression together provide independent levels of support for our identification of ASAPs as *bona fide* subunits of *T. gondii* F-type ATP synthase.

As can be expected for such an important protein complex playing a fundamental biological role, phylogenetic analysis revealed that most ASAPs identified from *T. gondii* are conserved across the phylum *Apicomplexa*. This suggests that the origin of a divergent F-type ATP synthase may predate the origin of apicomplexan parasites. In support of this conclusion, we find that many of the ASAPs are also conserved in two other alveolate phylums, *Chromerida* and *Dinoflagellata*. This opens up the opportunity for studying the structure and function of this unusual enzyme from free-living alveolate species, from which the enzyme can be isolated in native conditions at higher yields and purity. Moreover, a deeper understanding of the structure and function of this enzyme will facilitate the discovery of anti-parasitic compounds with pan-apicomplexan effect.

## Methods

### Sequence analysis and ortholog identification

#### Molecular reagents and methods

Genomic DNA and total RNA from tachyzoite stage *T. gondii*, and plasmid DNA from recombinant *E. coli* (DH5α) were isolated using Qiagen kits (Germany); cDNA was prepared from total RNA using the reverse transcription kit from Thermo Fisher Scientific (USA). The manufacturer’s protocol was followed while using the various kits. Polymerase chain reactions (PCR) was done using a proof reading DNA polymerase obtained from Takara Bio (Japan), and all primers used in this study (S1 Table) were obtained from Integrated DNA Technologies (USA). Ligase and various restriction endonucleases were purchased from New England BioLabs (USA) or Promega (USA).

#### Plasmid constructs

Various PCR products were cloned into the Topo 2.1 plasmid backbone (Thermo Fisher Scientific, USA) and sequenced before sub-cloning into other plasmids used for either modification of target genomic loci or ectopic expression of full-length cDNA for gene of interest in *T. gondii*. The plasmid construct used for tagging the 3’ end of F-type ATP synthase subunits β (*Tg*atpβ) and OSCP (*Tg*atpOSCP) with yellow fluorescent protein plus hemagglutinin (YFP-HA) tag had the following features: 2606 bp (BglII and AvrII) and 1831 bp (BamHI and AvrII) PCR fragments amplified from the 3’ end of *Tg*atpβ and *Tg*atpOSCP loci respectively, excluding the stop codon, were cloned into the Topo 2.1 plasmid, upstream of a YFP-HA tag coding sequence, followed by the 3’ UTR of the *T. gondii* dihydrofolate reductase-thymidilate synthase (*Tg*dhfr-ts) gene [58]. A modified pBluescript plasmid backbone was used for ectopic expression of full-length cDNA of selected *T. gondii* genes encoding the novel F-type ATP synthase subunits identified from this work. The cDNAs were cloned as BamHI and XbaI/NheI fragments in-frame to a HA tag, for constitutive expression using the *T. gondii* β-tubulin gene promoter and *Tg*dhfr-ts gene 3’ UTR. For isolating clonal lines of transgenic parasites, the respective plasmid constructs include either the DHFR cassette (for endogenous tagging; expresses a mutant version of the *Tg*dhfr cDNA that confers pyrimethamine resistance [59] or the CAT cassette (for ectopic expression; expresses the chloramphenicol acetyl transferase gene, which confers resistance to chloramphenicol [60].

#### T. gondii culture and genetic manipulation

Tachyzoite stage of *T. gondii* parasites (Type-I RH (RH) and Type-I RH ∆Ku80 [61] (RH∆Ku80)) strains were propagated following published protocols [62]. Briefly, Human foreskin fibroblast (HFF), which were used as host cells for parasite infection were grown in Dulbecco’s modified Eagle medium (DMEM High glucose) supplemented with 10% heat-inactivated fetal bovine serum, 2 mM GlutaMAX, 25 mM HEPES and 50 µg/ml Gentamicin. All cell culture grade reagents were procured from Thermo Fisher Scientific, USA. The HFF cells were maintained at 37°C in a humidified atmosphere containing 5% CO_2_. *T. gondii* tachyzoites were propagated by inoculating ~10^5^ freshly isolated tachyzoites into T25 flasks containing two-weeks old confluent HFF cell monolayer. The culture medium used for parasite growth is similar to that used for host cell growth except that it lacks serum. Parasites are harvested 48 h after infection by scrapping the host cell monolayer, physically disrupting the HFF cell suspension by passing it repeatedly through a 22-gauge needle, and filtering it through a 3 µM nucleopore membrane (Whatman, GE Healthcare, USA (to obtain a suspension of parasites devoid of HFF cell debris. The number of parasites present in 1 ml of suspension is estimated from cell counts obtained using a hemocytometer.

For generating transgenic parasites expressing the *Tg*ATPβ and *Tg*ATPOSCP proteins from the endogenous loci, the RH∆Ku80 parasites were transfected [61] with the respective tagging plasmids. Transfection was achieved by electroporating 10^7^ freshly harvested tachyzoite stage parasites resuspended in 400 µl parasite culture medium containing 50 µg of linearized sterile plasmid DNA, using a BioRad (USA) Gene Pulser system using 10µF capacitance, ∞ ohms resistance and 1.5 kV voltage settings. Transfected parasites were immediately inoculated into a T25 flask containing HFF monolayer and allowed to invade and replicated for 12-15 hours before beginning drug selection with 1 µM pyrimethamine. For ectopic expression of cDNA with HA tags for selected genes, RH parasites were transfected with the respective plasmid constructs and stable transformants were selected using 20 µM chloramphenicol. Clonal lines of stable transgenic parasites were isolated using the limiting dilution technique [62].

#### Measurement of intracellular ATP

For this assay, freshly isolated extracellular tachyzoite stage parasites were incubated in culture media which either contained 5.5 mM glucose and 4 mM glutamine or 0 mM glucose and 4 mM glutamine, in order to evaluate the ability of the parasites to maintain ATP homeostasis. The anti-parasitic drug atovaquone [43], an inhibitor of mtETC, was used to specifically inhibit mitochondrial ATP synthesis. After 2 h of incubation, total cellular ATP was measured using the ViaLight Plus Cell Proliferation and Cytotoxicity Bioassay Kit (Lonza, Switzerland) as per the manufacture protocol. The luminescence readout was quantified using the Varioskan Flash plate reader (Thermo Fisher Scientific, USA).

#### Microscopy and Western blotting

For visualizing the expression and subcellular localization of selected ASAPs fused to a reporter tag, replicating intracellular tachyzoite stage parasites stably expressing these proteins were imaged by microscopy. Before imaging the parasite infected HFF monolayers cultured on glass coverslips were washed with 1X phosphate buffered saline (PBS), and fixed using 4% paraformaldehyde for 20 min, before again washing and mounting on glass slides using the fluoroshield reagent (Sigma, USA). To counterstain the parasite mitochondrion, the cells were treated with 250 nM Mitotracker Red (Thermo Fisher Scientific, USA) for 20 minutes before fixing. The slides were imaged using the 63X oil immersion objective fitted to the Axio Observer inverted fluorescent microscope (Carl Zeiss, Germany), and images were processed using the Zen software (Carl Zeiss, Germany). Intrinsic fluorescence from YFP and Mitotracker Red were visualized using the excitation/emission filter combination of 493/520 and 578/599 respectively. HA tagged proteins were visualized by immunofluorescence staining, using rabbit α-HA primary antibodies (1:1000) followed by Alexa 488 conjugated goat anti rabbit secondary antibody (1:1000), both purchased from Thermo Fisher Scientific, USA. After fixing, the cells were permeabilized for 5-10 minutes using 0.25% Triton X-100 in 1X PBS. They were then treated with 2% fetal bovine serum in 1X PBS for 30 minutes, followed by primary antibody for 1 h, 3X wash with 1X PBS containing 0.25% Triton X-100, and secondary antibody for 1 h. Finally, the cells were washed 3X with 1X PBS containing 0.25% Triton X-100, followed by a water wash, before mounting the cover slip on glass slides for imagining.

For Western blotting, purified mitochondria were solubilized in 1X Laemmli buffer (120 mM Tris-HCL pH 6.8, 2% SDS, 10% glycerol and 0.01% w/v bromophenol blue) and resolved by SDS-PAGE, followed by electro-transfer to PVDF membrane (using Tris-glycine buffer pH 8 containing 20% methanol) at 65 mA for ~1 h 20 min. The blots were kept in blocking buffer (5% skimmed milk in 1X PBS) overnight at 4°C and then treated with primary antibody (1:5000; rabbit α-HA monoclonal antibody, Thermo Fisher Scientific, USA) for 1 h at ambient temperature. This was followed by 3X wash with 1X PBS containing 0.1% Tween 20, before incubating with horse radish peroxidase coupled secondary antibody (1:5000; donkey anti-rabbit antibody, Nif 824 GE Healthcare, USA) for 1 h at ambient temperature, and 3X wash with 1X PBS containing 0.1% Tween 20. The blot was then developed using either the 3,3’-diaminobenzidine substrate (Sigma, USA) or the chemiluminescent ECL Western blotting kit (GE Healthcare, USA).

#### Preparation of mitochondria from T. gondii

Freshly harvested tachyzoite stage transgenic parasites (~10^9^), expressing either *Tg*ATPβ-YFP-ΗΑ or *Tg*ATPOSCP-YFP-HAΗΑ, were washed with PBS and resuspended in 1-2 ml of hypotonic lysis buffer (15 mM phosphate buffer pH 7.5 and 2 mM Glucose). The cells were lysed by sonication in an iced water bath for 30 minutes, and samples were centrifuged at 2000 g for 2 minutes at 4°C to remove unbroken cells and large cell debris. The supernatant was recovered into a new tube and centrifuged at 21,000 g for 15 minutes at 4°C to pellet the mitochondria, which was then re-suspended in 500 µl of mitochondria storage buffer (320 mM sucrose, 1 mM EDTA, 10 mM Tris pH 7.4). Total protein in the mitochondria suspension was estimated by Bradford method [63] and the lysate was stored in −80°C until further use. When required, the mitochondria were recovered from storage buffer by centrifuging at 21,000 g for 15 minutes at 4°C and lysed using mitochondria solubilization buffer A (MSB-A; 50 mM NaCl, 50 mM Imidazole, 2 mM 6-Aminohexoanic acid, 1 mM EDTA, pH 7.0) containing β-dodecyl maltoside (DDM) detergent in total protein to detergent ratio of 1:5. Solubilization was allowed to continue overnight with rocking at 4°C, and the solubilized fraction was separated by centrifuging at 100,000 g for 20 minutes at 4°C.

#### Blue Native Page separation, in-gel ATPase activity assay and Western blotting

In order to identify the monomeric and dimeric forms of the *T. gondii* F-type ATP synthase, the solubilized mitochondrial lysate was subject to one-dimensional Blue Native PAGE (BNP) analysis [64,65]. Samples were prepared by adding glycerol to a final concentration of 5% and Coomassie Blue G 250 5% (w/v) dye was added such that DDM detergent to dye ratio was 8:1. Samples were separated on a 3-12 % native PAGE (Thermo Fisher Scientific, USA) at 150 V setting for 30 minutes in cathode buffer A, which was then switched to cathode buffer B and separation was continued till the blue dye exited the gel. The gel was visualized under white light to identify the various dye stained bands. 25 mM imidazole pH 7.0 was used as the anode buffer, while the composition of cathode buffers A and B were 50 mM Tricine, 7.5 mM imidazole pH 7.0, 0.02% Coomassie Blue G 250 dye and 50 mM Tricine, 7.5 mM imidazole pH 7.0, 0.002% Coomassie Blue G 250 dye, respectively.

For in-gel ATPase assay, after sample separation by BNP, the gel was incubated overnight in assay buffer (35 mM Tris-HCL pH 7.8, 270 mM glycine, 14 mM MgSO4, 0.2% Pb(NO_3_)_2_, and 8 mM ATP). The gel was then washed with water and the ATPase activity of F-type ATP synthase was observed by the formation of a milky white precipitate, visible against a black background, at the region corresponding to the expected monomeric and dimeric protein bands. For Western blotting after BNP separation, proteins were electro-transferred onto a PVDF membrane using Tris-glycine buffer pH 8.0 without methanol at 65 mAh for ~1hour 20 minutes. The membrane was then fixed in 8 % acetic acid for 15 minutes, air dried at room temperature for 30 minutes, and washed with methanol several times to remove the Coomassie dye stain [26]. Further steps were similar to that described above for SDS PAGE Western blotting.

#### Immunoprecipitation of F-type ATP synthase complex

The supernatant from solubilized mitochondria preparation from tachyzoite stage transgenic *T. gondii* expressing YFP-HA tagged *Tg*ATPOSCP subunit was used for immunoprecipitating the F-type ATP synthase using α-HA antibodies (Thermo Fisher Scientific, USA). The antibodies were first cross-linked to Protein A/G Plus agarose beads (Santa Cruz Biotechnology, USA) using the following protocol. ~50 µl of Protein A/G Plus agarose beads were washed twice in 500 µl 1X PBS (pH 7.4) at 4°C and mixed with 1 ml of 1X PBS containing 5-7 µg of rabbit α-HA antibody and incubate overnight at 4°C with gentle mixing on rocker. The beads were then separated by centrifugation (1000g for 2 minute at 4°C), equilibrate with 0.2 M Triethanolamine (pH 8.2) for 2 minutes and then washed twice with the same buffer. The bound antibodies were then cross-linked to the beads using 20 mM DMP (Sigma, USA) in 1 ml of 0.2 M Triethanolamine at room temperature for 45 minutes, followed by washing with 1ml of 50 mM Tris pH 7.5 twice for 15 minutes to quench the cross-linking reaction. Unbound antibodies are removed using three quick washes with 0.2 M glycine buffer pH 2.3. The antibody conjugated beads were then washed twice with 1X PBS and equilibrate with MSB containing the detergent DDM at critical micelle concentration. The supernatant from solubilized mitochondria was added to the antibody coupled beads and incubated overnight at 4°C with gentle mixing on rocker. The beads were then washed three times with 1X PBS containing DDM at critical micelle concentration. Proteins captured by the antibodies were eluted in 100 µl of 0.2 M glycine buffer (pH 2.3) in three rounds, and the pooled eluate was neutralized with 1M Tris (pH 8.0) and concentrated using a 3 KDa cut-off concentrator (Amicon Millipore, Germany). The samples were equilibrated with 0.1% RapiGest (Waters, USA) in 50 mM ammonium bicarbonate buffer in preparation for LC-MS analysis.

#### Purification of native F-type ATP synthase by chromatography

We followed a protocol that was previously reported for the purification of the F-type ATP synthase from the *Polytomella sp*. [66]. About 13 mg (wet weight) of mitochondrial preparation from transgenic *T. gondii* expressing YFP-HA tagged *Tg*atpβ subunit was solubilized overnight in 3 ml of mitochondrial solubilization buffer B (MSB-B; 50 mM Tris HCl (pH 8.0), 1 mM MgCl_2_, and DDM at protein to detergent ration 1:5). The supernatant was loaded onto a HiTrap DEAE sepharose column (GE Healthcare Life sciences, USA) having a bed volume of 1 ml and equilibrated with buffer A (50 mM Tris HCl (pH 8.0) MgCl_2_ 1mM, 0.05 % (w/v) DDM). After completion of loading, the column was first washed with 10 column volumes of buffer B_20_ (50 mM Tris HCl (pH 8.0), 1 mM MgCl_2_, 20 mM NaCl, and 0.05 % (w/v) DDM) and eluted with a 10X column volume linear gradient using buffer B_20_ and buffer B_500_ (50 mM Tris HCl (pH 8.0), 1 mM MgCl_2_, 500 mM NaCl, and 0.05 % (w/v) DDM) at a flow rate of 0.5 ml per minute. 500 µl fractions were collected and checked by Western blotting using α-HA antibodies. Peak positive fractions were identified, pooled and concentrated to a final volume of 200 µl using the Vivaspin concentrator (100 kDa cut-off; GE Healthcare Life sciences, USA) and separated by size exclusion chromatography using the Superose 6 Increase 3.2/300 column (GE Healthcare Life sciences, USA) having a bed volume of 2.4 ml. 50 µl sample was loaded on the column previously equilibrated with buffer B_20_, and then eluted with 1.5 column volumes of the same buffer at 50 µl flow rate. 100 µl fractions were collected and positive fractions, identified by Western blotting using α-HA antibodies, were pooled and concentrated using a 3 KDa cut-off concentrator (Amicon Millipore, Germany). Samples were then equilibrated with 0.1% RapiGest in 50 mM ammonium bicarbonate buffer in preparation for LC-MS analysis. All chromatography steps were carried out using the AKTApure system (GE Healthcare Life sciences, USA).

#### LC-MS proteomics

In order to identify the subunit components of the F-type ATP synthase from *T. gondii*, we performed LC-MS analysis on the partially purified enzyme obtained from BNP, immunoprecipitation and chromatographic purification. Sample preparation for LC-MS analysis was based on a previous report [67].

##### Sample preparation after BNP separation

The bands corresponding to the dimer and monomer forms of the protein were excised out from BNP gel, cut to small pieces, destained with 50% acetonitrile in 50 mM ammonium bicarbonate, and dehydrate with 100 % acetonitrile. The gel pieces were then treated with 10 mM DTT in 50 mM ammonium bicarbonate for 45-60 minutes at 56ºC to reduce the proteins, followed by alkylation in dark with 55 mM Iodoacetamide in 50 mM ammonium bicarbonate at ambient temperature for 45 minutes. Then, trypsinization of protein was carried out overnight at 37°C, and peptides were extracted using 50% acetonitrile in 2% formic acid. The extracts were dried using a speedvac and the peptides were reconstituted in 50 µl of 50 mM ammonium bicarbonate, acidified with HCl and desalted using the C_18_ Zip tip columns (Millipore, Germany). The samples were then dried in a speedvac and stored at −80°C until further use.

##### Sample preparation after Immunoprecipitation and size exclusion chromatography

Samples obtained from immunoprecipitation and size exclusion chromatography were immediately equilibrated with 0.1 % RapiGest as stated above. The mixtures were then heated to 80°C and maintained at this temperature for 15 minutes. The denatured proteins were reduced by 100 mM DTT at 60°C for 15 minutes, followed by alkylation with 200 mM iodoacetamide at ambient temperature and in dark for 30 minutes. Samples were then treated overnight with trypsin (Sigma Aldrich, USA) at 20:1 substrate to enzyme ratio at 37°C. Trypsinization was stopped by addition of 2 µl of 1N HCL and incubating at 37°C for 20 minutes, after which they were vortexed and centrifuged. The peptide samples were desalted using the C_18_ Zip tip (Millipore, Germany), dried and stored at −80°C until further use.

##### LC-MS analysis

The peptide samples were analyzed using LC-MS^E^ workflow on the nano ACQUITY (UPLC) Synapt system (Waters Corporation, USA). 4 µl of the digested peptide sample was injected into a 5 µm Symmetry C_18_ trapping column (180 µm x 20 mm) at a flow rate of 5 nl/min and peptides were eluted by using the following protocol: 3-40 % B (B - 100% acetonitrile with 0.1% formic acid) for 90 minutes, 40-85% B from 90-105 minutes and 97% A (A - 0.1% formic acid in water) and 3% B from 105-120 minutes, on a bridge ethyl hybrid C_18_ column (75 µm x 250 mm, 1.7 µm) at flow rate of 250 nl/min. The system was coupled to the Synapt High Definition Mass Spectrometer (Waters Corporation, USA) with a nonoLock spray as the ion source. Standard glu-fibrinopeptide B (Sigma Aldrich, USA) as the lockmass calibrate peptide was infused into ion source having 500 nl/min flow rate and sampled every 30 second. Acquisition of LC-MS^E^ data was performed in positive V mode with mass range m/z 50-1900 having scan time of 0.75 second, a constant low energy of 4 V for MS mode, and 20-40 V of collision energy during high energy MS^E^ mode scan. A capillary voltage of 3.5 kV and cone voltage 38 V were maintained during analysis. LC-MS^E^ data were processed using Protein Lynx Global Server (PLGS Version 2.5.3, Waters Corporation, USA) for identifying the proteins, with reference to the *T. gondii* proteome taken from *ToxoDB.org* release 36.

#### Gene co-expression analysis and phylogenetic studies

Normalized signal intensity values from microarray hybridizations performed at 13 different time points in the tachyzoite stage [51] were obtained directly from *EupathDB.org*. Normalized microarray intensities from time series data for intraerythrocytic stage *P. falciparum* were retrieved from reference [52]. Pearson correlation coefficients were calculated for all gene pairs using R (*R-project.org*). P-values were calculated using the Mann-Whitney U test [68] for co-expression between ATP synthase genes vs. co-expression between ATP synthase genes and non-ATP synthase genes, and for co-expression between ATP synthase genes vs. co-expression between non-ATP synthase genes. Since minute differences in very large samples may lead to very small p-values, the effect size was therefore tested using Cohen’s d test [69], where values above 0.8 denote large effect size. Orthologs for the novel subunits of *T. gondii* ATP synthase were identified as reported previously [41]. Sequences were aligned with mafft (v7.222) [70] and Neighbor-Joining trees were constructed using Clustalw (2.1) [71] using 1000 bootstrap replicates. The expected ‘true’ phylogeny was adopted from a previous study [41].

## Acknowledgements

The project is supported by funding from Science & Engineering Research Board (SERB), India (EMR/2016/003588), awarded jointly to DS and MB and This publication is also based upon work supported by the King Abdullah University of Science and Technology (KAUST) Office of Sponsored Research (OSR) under Award No. OCRF-2014-CRG3-2267 and the faculty baseline funding (BAS/1/1020-01-01) awarded to AP. RS received fellowship from Council of Scientific and Industrial Research, India. Authors would like to thank Dr. Mahesh J. Kulkarni and Ms. Rajeshwari Rathore, for LC-MS^E^ data acquisition and analysis, for Protein chromatography - Dr. Subashchandrabose Chinnathambi and Ms. Shweta Sonawane, CSIR-National Chemical Laboratory, Pune, and Prof. David. S. Roos, University of Pennsylvania, Philadelphia, USA, for providing *T. gondii* strains and other biological materials as gifts.

## Author contributions

DS conceived the project and designed the experiments along with RS, MB and AP. RS and TM executed the various experiments. DS, RS, AP and TM were involved in manuscript preparation.

